# CoolBox: a interactive genomic data explorer for Jupyter Notebook

**DOI:** 10.1101/614222

**Authors:** Weize Xu, Da Lin, Ping Hong, Liang Yi, Rohit Tyagi, Guoliang Li, Gang Cao

**Author notes:** To whom correspondence should be addressed, Contact: G.L. or G.C.

## Abstract

**Summary:** CoolBox is a Python package for interactive genomic data exploration based on Jupyter notebook. It provides a ggplot2-like Application Programming Interface (API) for genomic data visualization, and a Jupyter/ipywidgets based Graphical User Interface (GUI) for interactive data exploration. CoolBox is a versatile multi-omics explorer supporting most types of data formats generated by various sequencing technologies like RNA-Seq, ChIP-Seq, ChIA-PET and Hi-C.

**Availability and implementation:** CoolBox is purely implemented with Python, and the GUI widget in Jupyter notebook is based on the ipywidgets package. It is open-source and available under GPLv3 license at https://github.com/GangCaoLab/CoolBox.

## 1 Introduction

Multi-omics sequencing methods have greatly facilitated the functional genome research, such as RNA-Seq for transcriptome (Morin *et al.*, 2008), ChIP-Seq for DNA binding profile (Robertson *et al.*, 2007), ATAC-Seq (Buenrostro *et al.*, 2013) for chromatin accessibility, Hi-C (Lieberman-Aiden *et al.*, 2009) and ChIA-PET (Fullwood and Ruan, 2009) for chromatin organization. Integrative visualization of multi-omics data can greatly facilitate genome research, such as exploring the structure and behavior of the genome, finding featured patterns for further research. Several tools have been developed for this purpose, such as WashU Epigenome Browser (Zhou *et al.*, 2011) and the 3D genome browser (Wang *et al.*, 2018). These kinds of browsers implemented as web service can be accessed over Internet with a web browser and are easy for biologists without any computational background to use. However, they can not be manipulated conveniently by programming. Moreover, the connection between the client and server over the internet might be often slow. Another kind of tool is the command line plotting tool such as HiCPlotter (Akdemir and Chin, 2015) and pyGenomeTracks (Ramírez *et al.*, 2018). They can plot the figures in a programming way and can be run on local machine. Yet it cannot provide the GUI for interactive visualization. Thus, a new genomic data visualization tool which combines advantages of both kinds of tools to provide a GUI for interactive visualization, an API for programing control and can be run on a regular PC is desperately wanted for bioinformaticians and data scientists.

Jupyter Notebook (http://jupyter.org/) is an open-source and highly interactive data analysis platform. It allows users to write documents together with executable code, rich text and media. The interactive Graphical User Interface (GUI) can be created within the document through the ipywidgets package. Here, we developed CoolBox, which implemented a Python API for genomic data visualization and a simple GUI for data exploration within Jupyter notebook environment. It helps users to compose and configure the figure or browser object in a flexible and programmable way. Meanwhile, CoolBox provides an interface for fetching precise data, facilitating the graphic visualization and statistical analysis. Through embedding on the Jupyter notebook, CoolBox is more convenient to showcase the visualization results with high reproducibility and integrate with other Python packages.

## 2 CoolBox

### 2.1 Interactive genomic data exploration

As shown in Fig. 1, CoolBox supports the interactive data visualization, by which users can explore different genomic regions by operating a simple widget panel and visualize the data within this region. In addition, the data and the figures are bound together by Python objects. In this way, the users can get the precise data of each track within the current view of the genome region through the API. Such design facilitates the comparative visualization and statistical analysis.

**Fig. 1.**
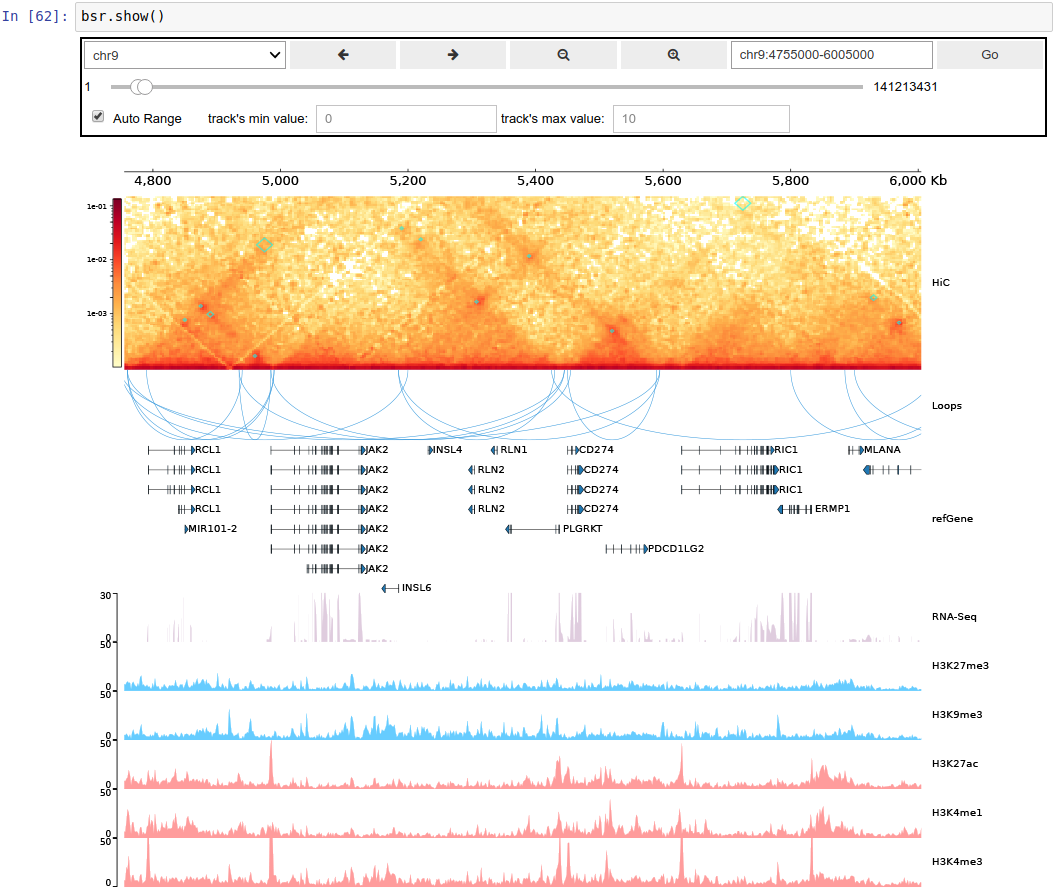
A CoolBox example in the Jupyter notebook. A browser object is created by code as shown in the highlighted box. When the “.show()” API is called, a simple widget and the plotted tracks will be shown. Users can explore different genome regions interactively by using the widget.

Most sets of commonly generated genomic assay data such as, ChIP-Seq data, Hi-C data, ATAC-Seq data etc. can be shown in CoolBox by different kinds of tracks. Most tracks’ features (color, height, style etc.) can be configured exactly via the API. In CoolBox plotting system, the plots are not only single layer – users can put another layer (Coverage) upon the original plot to produce more comprehensive figures. Moreover, the output figures can be conveniently saved to different kinds of image formats such as PNG, JPEG, PDF and SVG.

### 2.2 API design

The API design of CoolBox is inspired by popular R package ggplot2 (Wickham, 2010). Users can use the “+” operator to integrate the low-level elements together to obtain the higher-level elements, like composing track objects to frame or browser object. Through this API design, a browser object can be constructed by a Python expression (as the code shown in Fig.1). More details about the API are available in online documents.

### 2.3 Implementation

The plotting system of CoolBox is based on the matplotlib package. A part of plotting code in the CoolBox is transplanted from pyGenomeTracks package (Ramírez *et al.*, 2018). The data loading of bigWig and “.cool” file format are used from the pyBigWig (Ryan *et al.*, 2016) and cooler package (Abdennur *et al.*, 2018). The GUI is based on the ipywidgets package.

### 2.4 Example usage

We have performed many visual data exploration using CoolBox on our unpublish multi-omics data from THP-1 cell MTB infection experiments. An example is shown in Fig. 1, within this genome region we can see many gene’s promoter is located at TAD’s boundary, and highly expressed genes have a relatively higher activate histone modification in their promoter region. User can use CoolBox to perform similar visualization analysis on their own dataset, more detail are available in our online documents.

## 3 Conclusion

CoolBox is a versatile toolkit for the visualization of most types of genomic data and exploring multi-omics data in Jupyter notebook. It provides a user-friendly ggplot2-like Python API, a GUI for browsing the data, and a convenient tool to analyze the figures and their corresponding data simultaneously. Through the power of Jupyter notebook, it provides an easy way for bioinformaticians to exploit this tool’s versatility for better data demonstration. Of note, the code and figures can be organized into the same page, which could increase the reproducibility of genomic data visualization tasks.

## Acknowledgements

The authors would like to thank the pyGenomeTracks project development team for their open source plotting code, which is very useful for building the CoolBox.

## Funding

This work was supported by the National Key Research and Development Project of China (Grant No.2017YFD0500303, 2016YFA0102500), the National Natural Science Foundation of China (Grant No.31771402, 91440114) and Huazhong Agricultural University Scientific & Technological Self-innovation Foundation (Program No.510318177).

